# Can Neuropathic Pain Predict Response to Arthroplasty in Knee Osteoarthritis? A Prospective Observational Cohort Study

**DOI:** 10.1101/360412

**Authors:** Anushka Soni, Kirsten M Leyland, Amit Kiran, Nigel K Arden, Cyrus Cooper, Vishvarani Wanigasekera, Irene Tracey, M Kassim Javaid, Andrew J Price

**Author notes:** **Corresponding author:** Dr. Anushka Soni, Wellcome Centre for Integrative Neuroimaging, FMRIB, Nuffield Department of Clinical Neurosciences, University of Oxford, Oxford, OX3 9DU.

## Abstract

A significant proportion of patients with knee osteoarthritis (OA) continue to have severe ongoing pain following knee replacement surgery. Central sensitization and features suggestive of neuropathic pain before surgery may result in a poor outcome post-operatively. In this prospective observational study of patients undergoing primary knee arthroplasty (n=120), the modified PainDETECT score was used to divide patients, with primary knee OA, into nociceptive (<13), unclear (13–18) and neuropathic -like pain (>18) groups pre-operatively. Response to surgery was compared between groups using the Oxford Knee Score (OKS) and the presence of moderate to severe long-term pain 12 months after arthroplasty. The analyses were replicated in a larger independent cohort study (n=404). 120 patients were recruited to the main study cohort: 63 (52%) nociceptive pain; 32 (27%) unclear pain; 25 (21%) neuropathic-like pain. Patients with neuropathic-like pain had significantly worse OKS pre and post-operatively, compared to the nociceptive pain group, independent of age, sex and BMI. At 12-months post-operatively the mean OKS was 4 points lower in the neuropathic-like group compared with the nociceptive group in the study cohort (non-significant); with a difference of 5 points in the replication cohort (p<0.001). Moderate to severe long-term pain after arthroplasty at 12-months was present in 50% of the neuropathic-like pain group versus 24% in the nociceptive pain group, in the replication cohort (p<0.001). Neuropathic pain is common and targeted therapy pre, peri and post-operatively may improve treatment response.

## INTRODUCTION

Knee Osteoarthritis (OA) is a common chronic disease which typically presents with disabling joint pain, stiffness and loss of function, and progresses with age [2; 23; 36]. In the UK, the estimated prevalence of symptomatic knee OA is 11% to 19% [44; 48] and the lifetime risk of total knee replacement surgery is 10% for males and 8% for females in those aged over 50 [14]. Whilst knee replacement surgery is widely regarded as an effective intervention, 15% of patients report severe-extreme persistent pain post-operatively [8; 63].

In England, Wales, Northern Ireland and Isle of Man, 108,713 knee replacement procedures were performed in 2016 and, with the aging population, an exponential rise in demand for costly surgery is predicted [13; 14; 16; 33] [62]. Identifying patients pre-operatively who are not likely to improve post-surgery is essential for optimal management, the selection of patients, and efficient service delivery.

Indeed, the strongest independent predictors of persistent pain after TKA are pre-operative pain catastrophizing and knee pain severity [30]. These findings emphasize the importance of evaluating the multidimensional nature of pain and it may be that factors operating beyond the constraint of the primary affected joint itself should be considered.

Furthermore, there is increasing evidence to support the presence of central sensitization and features of neuropathic pain in a subgroup of patients with osteoarthritis (OA) [4; 11; 17-19; 24-26; 37; 45; 47; 54; 60]. It has been proposed that understanding the altered central pain modulation may represent an opportunity to predict response to treatments, as well as enabling more targeted therapy in OA [1; 3; 5; 37; 59].

However, features suggestive of neuropathic pain are not currently routinely assessed when patients are being considered for knee replacement surgery. The two main tools available for potential use in clinical practice are quantitative sensory testing (QST) and screening tools such as the PainDETECT questionnaire (PD-Q). PD-Q has several benefits as a validated and simpler screening tool, and it has specifically been modified for use in knee OA [7; 12; 26; 59]. Identifying altered pain processing preoperatively, with QST, has been suggested to help predict the development of persistent post-operative pain, but this is not yet part of routine clinical practice [38; 39; 49; 51; 64].

In this study, we aimed to describe the prevalence of neuropathic-like pain features, using the PD-Q, in a cohort of patients with knee OA awaiting primary knee replacement surgery, and to compare the patient reported outcomes in patients identified pre-operatively as having nociceptive, unclear, or neuropathic-like pain. Our hypothesis was that the patients with features of neuropathic pain would have worse outcomes following surgery even after accounting for difference in severity of symptoms prior to surgery.

## PATIENTS AND METHODS

We primarily used data from the Evaluation of Peri-operative pain In Osteoarthritis of the kNEe (EPIONE) Study (baseline, 2months and 12months post-surgery) from a NHS single site; and then went on to replicate our overall findings (baseline and 12months) using a second independent patient cohort; Clinical Outcomes in Arthroplasty Study (COASt) from two large NHS hospitals.

### Study cohort: EPIONE Study

#### Setting and patient recruitment

The EPIONE Study is a prospective cohort study of patients with primary OA, awaiting primary knee replacement surgery who underwent detailed pain assessment prior to surgery. The study was conducted at the Nuffield Orthopaedic Centre, as part of Oxford University Hospitals NHS Trust, a specialist referral centre for joint replacement surgery. Patient recruitment took place at their routine pre-operative assessment clinic appointment within 6 weeks of surgery (baseline) between October 2011 and May 2014. Patients seen by the Consultant orthopaedic surgeon led outpatient Knee team at the Centre, were invited to participate if they were undergoing primary total or uni-compartmental arthroplasty for primary osteoarthritis of the knee. Patients were excluded if they were unwilling or unable to give informed consent; unable to complete questionnaires; had a poor understanding of English; known Charcot’s arthropathy or other severe neurological disorders; were undergoing revision arthroplasty, or had secondary OA due to trauma. A sub-group of patients recruited to the main study cohort were invited to participate in a neuro-imaging sub-study but these data will not be presented here [55].

#### Data collection

Data collection took place prior to surgery alongside the routine pre-operative assessment clinic appointment within 6 weeks of surgery, at 2 months postoperatively to coincide with the routine clinical follow-up appointment, and at 12 months post-operatively to capture mid-term outcome data [49]. The follow-up assessments were conducted by post. The participants who did not initially respond were sent two postal reminders. At baseline patients provided information on their age, sex, marital status, employment status, which knee was predominantly affected and the type of surgery planned. Further clinical data were collected from the hospital’s electronic patient clinical record system and included height, weight and medications prior to surgery. Weight-bearing anterio-posterior films were taken as part of routine clinical care and scored using the Kellgren and Lawrence global score [29; 30] by a single observer (KML) who was blinded to patient identity and symptoms. Radiographic severity was dichotomized using a Kellgren and Lawrence score of grade three or higher in order to identify subjects who at least had definite osteophytes and joint space narrowing [30].

##### Pain assessment (exposure groups)

At baseline, the quality of pain was assessed using the modified form of the PainDETECT questionnaire (mPD-Q) [7; 12; 26; 59], and the short form of the McGill Pain Questionnaire (SF-MPQ)[40; 41]. The mPD-Q score was used to divide patients, according to established cut-off values, into those with nociceptive (<13), unclear (13–18) and neuropathic-like pain (>18) [20] [22; 24; 26]. The SF-MPQ was used to rank intensity reported for sensory, affective and total descriptors of pain[40]. Participants also completed the following questionnaires: the Hospital Anxiety and Depression Scale (HADS) [9], the state/trait anxiety inventory [56], the pain catastrophising scale (PCS) [57], the Tampa scale for kinesiophobia (TSK) [32], the revised Life Orientation Test (LOT-R) [53] and the Pittsburgh Sleep Quality Index (PSQI) [10].

##### Primary outcome measure

The primary outcome measure was the difference in Oxford Knee Score (OKS) between the nociceptive and neuropathic pain groups. The OKS is a 12-item composite score developed in order to measure patient reported outcome after total knee replacement (TKR), which measures three symptom domains: pain, stiffness, and functional disability, in relation to the knee [15]. The latest recommendation is to use a scoring system which gives a summary score ranging from 0 (worst possible score, most severe symptoms) to 48 (best possible score, least symptoms) [42]. Patients completed the OKS at baseline, prior to surgery, and at 2 months and 12 months post-operatively. The participants who did not initially respond were sent two postal reminders. The study was designed in order to detect the minimal clinically important difference in the Oxford Knee Score (OKS) between the neuropathic and nociceptive pain groups. It was calculated that to an approximate sample size of 148 patients would be needed to detect a difference of 5 points on the OKS between the two pain groups, with a power of 80%, a type I error of 5% and an estimated prevalence of neuropathic pain of 25% [6] [25; 26],

##### Secondary outcome measures

Secondary outcome measures included the proportion of patients achieving a 7 point improvement in OKS from pre-surgical baseline, which is the minimally clinically important change in OKS, for use in individual patients, as well as the proportion of patients with moderate to severe long-term pain after arthroplasty in each pain group. [6]. Moderate to severe long-term pain after arthroplasty was defined as an average pain severity score of three or more for the preceding week 12 months after surgery measured using the visual analogue scale within the SF-MPQ which was repeated at 2 and 12 months post-operatively [49].

The local ethics committee approved the study and written consent was obtained from each participant (NRES Committee-South Central-Oxford B, 09/H0605/76).

### Replication cohort: COASt study

For the replication analysis, we used data from the Clinical Outcomes in Arthroplasty Study (COASt). In brief, eligible patients who were placed on the waiting list for knee or hip replacement surgery were recruited across two hospitals between 2010 and 2013 (baseline): Southampton University Hospital NHS Foundation Trust (UHS) and Nuffield Orthopaedic Centre (NOC) as part of Oxford University Hospitals NHS Trust. Patient demographics and clinical data were collected the pre-operative outpatient visit prior to surgery as well as annual post-operative follow-up by post for five years thereafter. In line with the EPIONE study, the cohort from COASt was restricted to the patients who were listed for knee replacement surgery and had 12 month follow up data available. Baseline data included age, sex, BMI, employment status, side predominantly affected, duration of symptoms and the type of surgery planned. Pain was assessed before and 12 months after surgery using the mPD-Q. OKS was also reported at baseline and at 12 months post-surgery. Medication use prior to surgery was not available, and so could not be included as a potential confounder. There was no overlap between participants in the study and replication cohorts.

### Statistical analysis

Summary statistics were used to describe the demographic and clinical baseline data in each of the three pain groups for EPIONE and COASt separately. Pre-operative demographic and clinical patient characteristics for the unclear and neuropathic pain groups were compared to the nociceptive group, using Student’s t-test, Wilcoxon-Mann-Whitney, and Chi-square test for normally distributed, non-normally distributed, and categorical data respectively.

The difference in Oxford Knee Score (OKS) between those with nociceptive, unclear and neuropathic pain was compared, pre-operatively and at 2 and 12-months postoperatively, using repeated measures generalized estimating equations (GEE) linear regression models. The potential confounding factors included in the model were age, sex and BMI and were selected a priori [35; 58]. Regression diagnostics checking for normality of residuals, collinearity, homoscedasticity, and linearity were conducted. Assumptions for the GEE regression models were met in all analyses. There was no missing data for the included confounding factors. The analyses were not restricted to patients who had completed assessments at all the time points such that in the case of missing data for the outcome variable, observations available for any other time-points were included in the model.

Logistic regression modelling was then used to test if there was any significant relationship between the moderate to severe long-term pain after arthroplasty and pain grouping. Pain severity was collected from the SF-MPQ in the study cohort. For COASt, pain was measured using the numeric pain rating average pain severity score, captured using the mPD-Q at the 12-month follow up assessment. The models were repeated with the definition of moderate to severe long-term pain after arthroplasty as a score of 4 and then 4.5 in order to assess the effect of a more stringent definition of moderate to severe long-term pain after arthroplasty.

All analyses were conducted using Stata SE v12.0 (StatCorp, College Station, TX, USA)

## RESULTS

The recruitment process and study visits for EPIONE are outlined in Figure 1.

**Figure 1:**
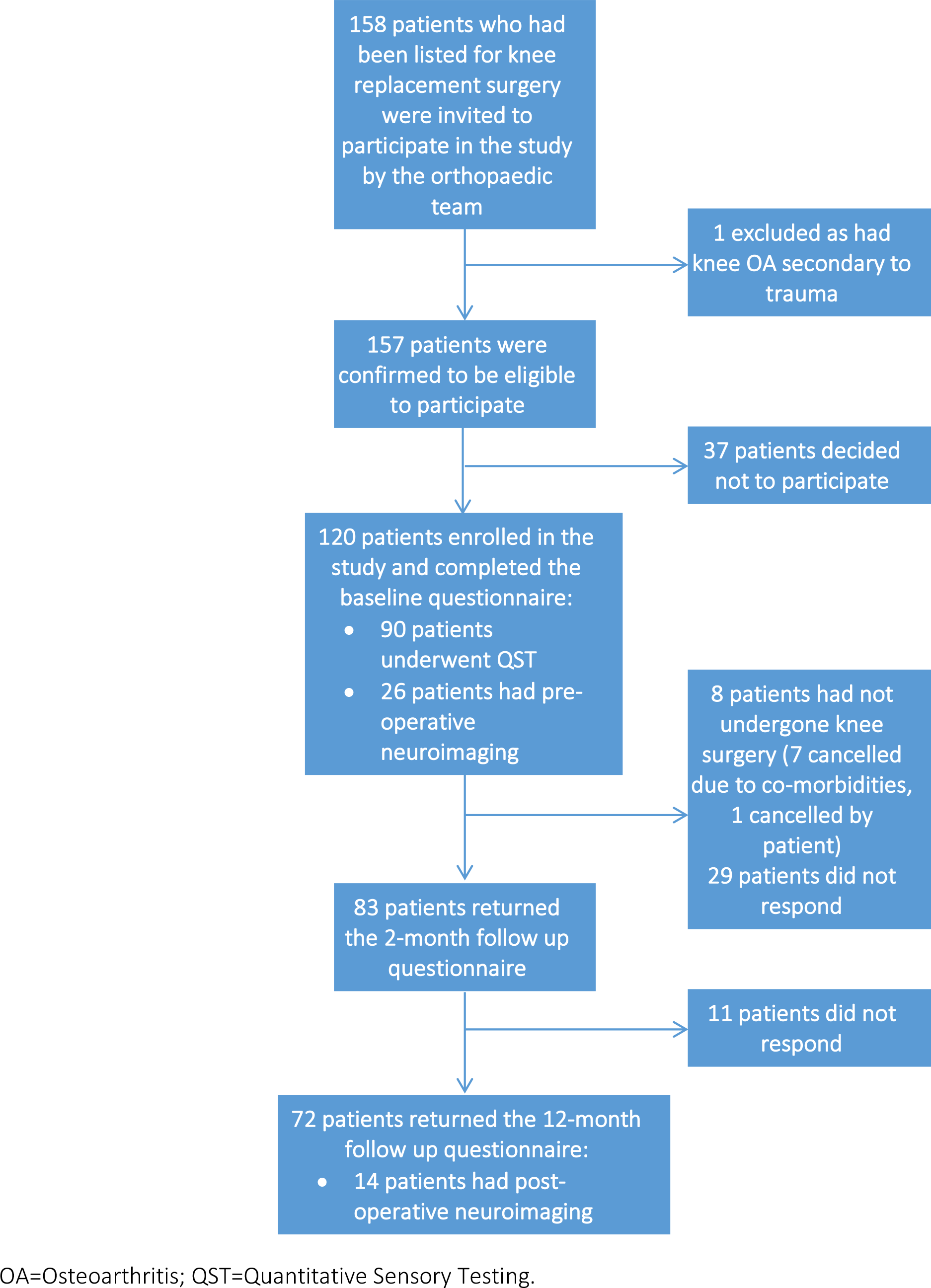
Diagram of study recruitment and follow up visits for the EPIONE study.

### The EPIONE Study

Among the 120 patients recruited to EPIONE, at baseline 25 (20.8%) had an mPD-Q score characterized as predominantly neuropathic pain, 32 (26.7%) had unclear pain, and 63 (52.5%) had predominantly nociceptive pain. Baseline demographics and clinical features (**table 1**) were similar across the three pain groups, except for age, SF-MPQ score, additional pain locations and K/L score. The neuropathic-like pain group were significantly worse than the nociceptive pain group across almost all psychological scales including anxiety, depression, catastrophizing and sleep quality.

**Table 1.** Pre-operative patient characteristics of the 120 patients recruited to EPIONE divided into nociceptive, unclear and neuropathic pain groups*.

|  | Nociceptive pain<br>(n=63) | Unclear pain<br>(n=32) | Neuropathic pain<br>(n=25) |
| --- | --- | --- | --- |
| **Demographic features** |  |  |  |
| Age, mean ± SD years | 72 (8) | 68 (8) <sup>†</sup> | 70 (10) |
| Female, n (%) | 27 (43) | 20 (63) | 14 (56) |
| BMI, mean ± SD kg/m<sup>2</sup> | 29.5 (5.1) | 30.2 (5.2) | 31.8 (4.9) |
| Employed, n (%) | 14 (22) | 12 (38) | 3 (13) |
| Married or living with partner, n (%) | 47 (75) | 18 (58) | 15 (60) |
| **Clinical features** |  |  |  |
| Right knee affected, n (%) | 35 (55) | 18 (56) | 10 (43) |
| Duration of pain, median (IQR) months | 36 (15-90) | 60 (24-120) | 48 (36-120) |
| Use of pain-modifying medication, n(%) | 36 (57) | 22 (69) | 17 (68) |
| SF-MPQ total score, mean ± SD range 0-45 | 16.3 (9.5) | 23.0 (9.4) <sup>†</sup> | 24.5 (7.7) <sup>††</sup> |
| Number of additional body areas which have caused pain for at least 3 months: |  |  |  |
| 0, n (%) | 19 (30) | 15(47) | 7 (28) |
| 1-2, n (%) | 27(42) | 5 (15) | 7 (28) |
| $\geq 3$, n (%) | 17 (27) | 12 (37) | 11 44) |
| Kellgren and Lawrence grade, n (%): |  |  |  |
| 0-2 | 4 (7.1) | 6 (20.7) | 4 (17.4) |
| 3-4 | 52 (92.9) | 23 (79.3) | 19 (82.6) |
| **Psychological characteristics** |  |  |  |
| HAD Anxiety, mean ± SD range 0-21 | 6.44 (4.2) | 7.2 (5.0) | 9.3 (4.0) <sup>†</sup> |
| HAD Depression, mean ± SD range 0-21 | 6.3 (3.3) | 7.3 (4.8) | 8.6 (4.2) <sup>†</sup> |
| STAI State anxiety, mean ± SD range 20-80 | 34.0 (12.5) | 39.7 (14.2) <sup>†</sup> | 41.0 (14.7) <sup>†</sup> |
| STAI Trait anxiety, mean ± SD range 20-80 | 33.4 (10.7) | 37.3 (12.9) | 43.2 (15.9) <sup>†</sup> |
| Pain Catastrophising Score, median (IQR) range 0-52 | 11 (6-17) | 19 (10-28) <sup>†</sup> | 21 (10-36) <sup>††</sup> |
| Life orientation Test-R, mean ± SD range 0-24 | 16.8 (4.3) | 15.4 (5.5) | 12.5 (5.9) <sup>††</sup> |
| Pittsburgh Sleep Quality Index, mean ± SD range 0-21** | 8.6 (3.3) | 10.0 (3.9) | 10.8 (4.0) <sup>†</sup> |
| Tampa scale of kinesophobia, mean ± SD range | 38.3 (9.8) | 39.7 (7.6) | 42.4 (4.9) |

\* The Pain-DETECT questionnaire was used to divide patients, according to established cutoff values, into those with nociceptive (<13), unclear (13–18) and neuropathic pain (>18). P-values were calculated in comparison to the nociceptive group, <sup>†</sup>p<0.05 <sup>††</sup>p<0.001; SD= standard deviation; IQR=interquartile range. \*\*Measures of Pittsburgh Sleep Quality Index were only available for 49, 23 and 20 participants in the nociceptive, unclear and neuropathic pain groups respectively.

### Replication: COASt cohort

In the replication analysis, 404 patients from the COASt study with pre-operative data available were included in this analysis. Of these 59 (15%) had neuropathic pain, 112 (28%) had unclear pain and 233 (58%) had nociceptive pain. The pre-operative characteristics for the participants were similar across all groups except for age, BMI and depression (table 2).

**Table 2.**
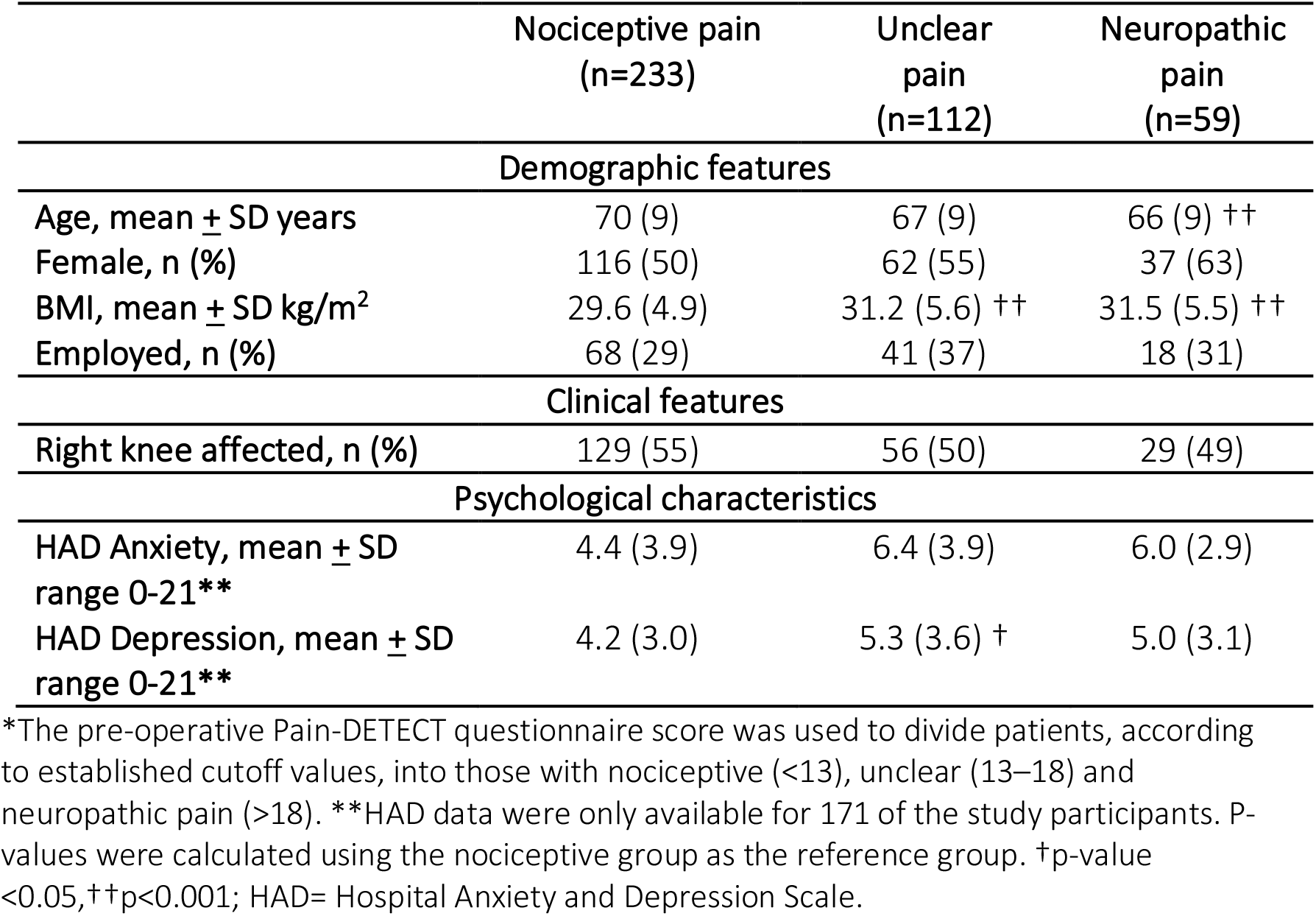
Pre-operative clinical, pain and psychological characteristics of 404 patients recruited to COASt, divided into nociceptive, unclear and neuropathic pain groups*.

|  | Nociceptive pain<br>(n=233) | Unclear<br>pain<br>(n=112) | Neuropathic<br>pain<br>(n=59) |
| --- | --- | --- | --- |
| **Demographic features** |  |  |  |
| Age, mean ± SD years | 70 (9) | 67 (9) | 66 (9) $\dagger\dagger$ |
| Female, n (%) | 116 (50) | 62 (55) | 37 (63) |
| BMI, mean ± SD kg/m<sup>2</sup> | 29.6 (4.9) | 31.2 (5.6) $\dagger\dagger$ | 31.5 (5.5) $\dagger\dagger$ |
| Employed, n (%) | 68 (29) | 41 (37) | 18 (31) |
| **Clinical features** |  |  |  |
| Right knee affected, n (%) | 129 (55) | 56 (50) | 29 (49) |
| **Psychological characteristics** |  |  |  |
| HAD Anxiety, mean ± SD<br>range 0-21** | 4.4 (3.9) | 6.4 (3.9) | 6.0 (2.9) |
| HAD Depression, mean ± SD<br>range 0-21** | 4.2 (3.0) | 5.3 (3.6) † | 5.0 (3.1) |

\*The pre-operative Pain-DETECT questionnaire score was used to divide patients, according to established cutoff values, into those with nociceptive (<13), unclear (13–18) and neuropathic pain (>18). \*\*HAD data were only available for 171 of the study participants. P-values were calculated using the nociceptive group as the reference group. † p-value <0.05, $\dagger\dagger$ p<0.001; HAD= Hospital Anxiety and Depression Scale.

### Descriptive data for study and validation cohorts

In EPIONE, the neuropathic pain group had significantly worse OKS scores (mean [95% CI]) compared to the nociceptive pain group, both pre-operatively (13.4 [10.2-16.6] versus 20.3 [18.3-22.3]) and 2-month post-operatively (29.2 [25.1-33.2] versus 35.8 [33.5-38.0]) in unadjusted and adjusted models (age, sex and BMI) (**table 3). Figure 2** shows that this same trend was present at 12-months post-operatively, but did not reach statistical significance (35.4 [31.3-39.4] versus 39.7 [37.4-42.0]). When pre-operative pain severity and pain-modifying medication use were included in the models, the OKS remained significantly lower in the neuropathic group compared to the nociceptive group prior to surgery (20.7 [17.9-23.4]) versus 25.8 [24.0-27.5]) and at 2-months post-operatively, (24.3 [21.1-27.4] versus 30.4 [28.5-32.3]). Similar associations were seen in the COASt cohort with significant findings both pre and post operatively (**Figure 2**).

**Table 3.** OKS scores for EPIONE and COASt before and after surgery.

| EPIONE | Nociceptive pain<br>(n=63)* | Unclear pain<br>(n=32)* | Neuropathic pain<br>(n=25)* |
| --- | --- | --- | --- |
| OKS Score<br>mean (SD); median (IQR) |  |  |  |
| Pre-op | 20.5 (7.3) | 19.2 (7.5) | 13.1 (5.5) |
| 2-months post-op | 39.0 (29.0-46.0) | 35.0 (29.0-40.0) | 32.0 (18.0-41.0) |
| 12-months post-op | 43.0 (38.0-46.0) | 43.5 (35.0-44.5) | 39.0 (32.0-43.0) |
| 7 point MCIC in OKS<br>(pre-op to 12-months n (%)) | 37/42 (88) | 15/16 (94) | 12/14 (85) |
| Long-term pain after<br>arthroplasty, n(%) | 6/42 (14) | 6/16 (38) | 5/14 (36) |
| COAST | Nociceptive pain<br>(n=233) † | Unclear pain<br>(n=112) † | Neuropathic pain<br>(n=59) † |
| OKS Score<br>mean (SD); median (IQR) |  |  |  |
| Pre-op | 22.3 (7.7) | 18.9 (6.8) | 15.5 (5.4) |
| 12 months post-op | 42.0 (35.0-46.0) | 39.0 (29.5-44.0) | 37.0 (25.0-43.0) |
| 7 point MCIC in OKS<br>(pre-op to 12-months n (%)) | 119/219 (91) | 89/107 (83) | 46/58 (79) |
| Long-term pain after<br>arthroplasty, n(%) | 53/219 (24) | 44/107 (41) | 29/58 (50) |

\*number at baseline; at 2-month visit there were 46, 23, and 14 patients in the nociceptive, unclear and neuropathic groups respectively. At 12-months there were 42, 16 and 14 patients in the nociceptive, unclear and neuropathic groups respectively. †number at baseline; at 12-month visit there were 219, 107, and 58 patients in the nociceptive, unclear and neuropathic groups respectively. OKS=Oxford Knee Score. MCIC=Minimally Clinically Important Change.

**Figure 2.**
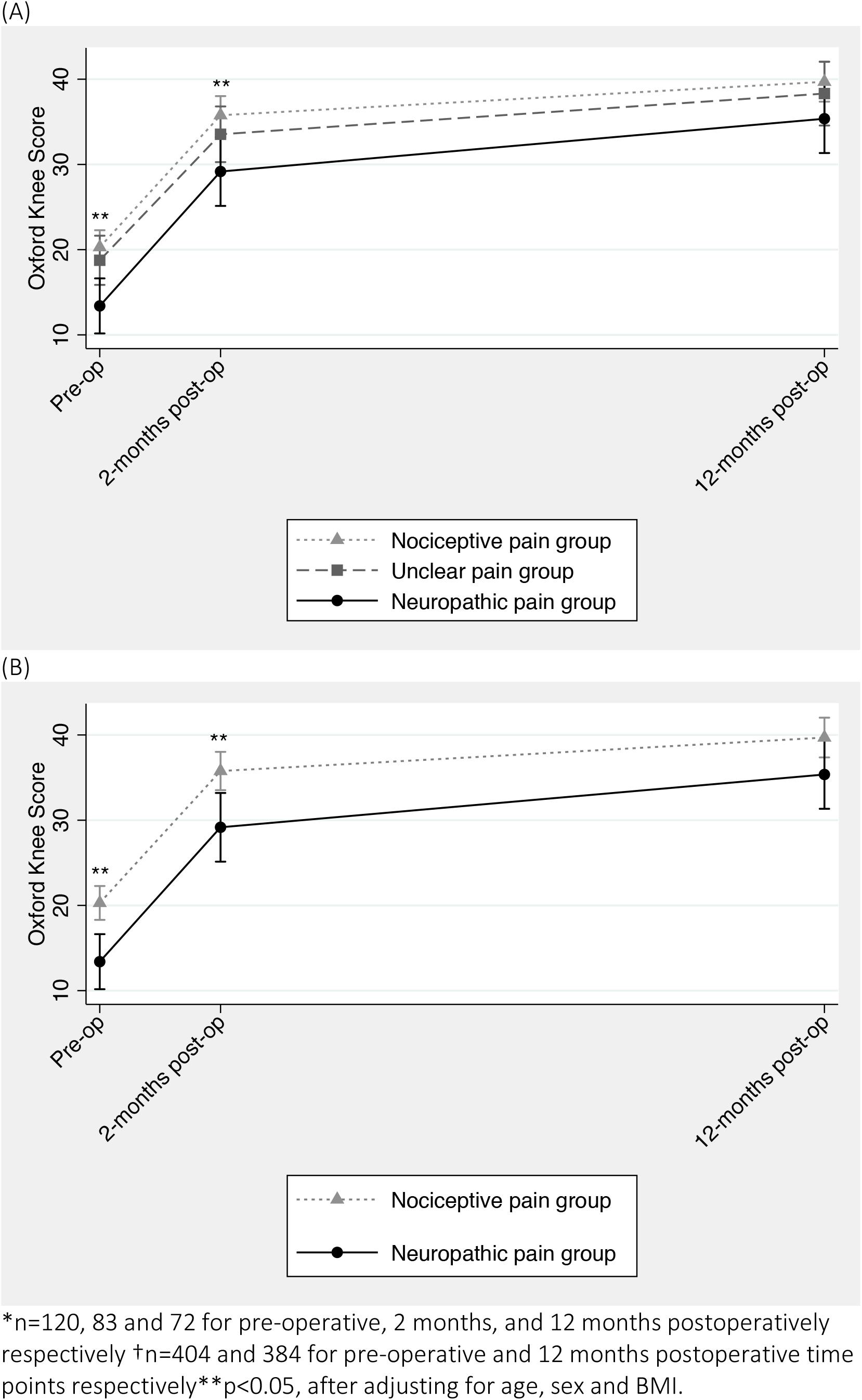
Oxford Knee Score for the participants of (A) EPIONE* and (B) COASt† for all pain subgroups.

#### Long-term pain after arthroplasty

In the EPIONE cohort 72 patients had follow-up data at 12months and 23% (17/72) of the patients met the criteria for moderate to severe long-term pain after arthroplasty at the 12-month follow up assessment. Of those with nociceptive pain prior to surgery, 6/43 (14%) reported moderate to severe long-term pain after arthroplasty compared to 6/16 (38%) in the unclear group, and 5/14 (36%) in the neuropathic pain group. Univariable logistic modelling showed that patients in the unclear pain group prior to surgery were significantly (p=0.05) more likely to report moderate to severe long-term pain after arthroplasty at 12-months post-operatively, compared to the nociceptive group (**table 4**). There was a similar, but non-significant trend, in the neuropathic pain group. However, when confounding factors, including age, sex, BMI and pain severity at baseline, were included in the model, there was no significant difference in the prevalence of moderate to severe long-term pain after arthroplasty between the pain groups or when a higher cut-off value was used to define moderate to severe long-term pain after arthroplasty (**Appendix**).

In the COASt cohort, 126 of the study participants met the criteria for moderate to severe long-term pain after arthroplasty. Of those with nociceptive pain prior to surgery, 24% (53/219) reported moderate to severe long-term pain after arthroplasty, compared to 41% (44/107) in the unclear group and 50% (29/58) in the neuropathic group. Patients in the unclear and neuropathic groups, as determined prior to surgery, were significantly more likely to report moderate to severe long-term pain after arthroplasty, when compared to the nociceptive group, **table 4**. This relationship remained significant after adjusting for the effects of age, sex, BMI and pain severity prior to surgery and when a higher threshold for moderate to severe long-term pain after arthroplasty was used (**Appendix**).

**Table 4.** Logistic regression model of the association between pain group at baseline and moderate to severe long-term pain after arthroplasty at 12-month follow up assessment.

| Predictor | Univariable model |  | Multivariable model* |  |
| --- | --- | --- | --- | --- |
|  | OR (95% CI) | p | OR (95% CI) | p |
| **EPIONE** |  |  |  |  |
| **Nociceptive group** | Reference group | - | Reference group | - |
| **Unclear group** | 3.7 (1.00-13.99) | 0.05 | 3.4 (0.87-13.77) | 0.08 |
| **Neuropathic group** | 3.4 (0.85-13.79) | 0.08 | 2.6 (0.55-12.43) | 0.23 |
| **COASt** |  |  |  |  |
| **Nociceptive group** | Reference group | - | Reference group | - |
| **Unclear group** | 2.3 (1.4-3.8) | 0.001 | 2.0 (1.2-3.4) | 0.006 |
| **Neuropathic group** | 3.6 (2.0-6.6) | <0.001 | 3.0 (1.6-5.8) | 0.001 |

\*Adjusted for age, sex, BMI and pain severity prior to surgery. OR=odds ratio; 95% CI=95% confidence interval.

## DISCUSSION

This is one of the first studies to examine the influence of pre-operative neuropathic pain in knee OA on outcomes of knee replacement surgery, the features of which are not routinely assessed when considering patients for knee replacement surgery. The key finding of this study is that patients with features of neuropathic pain, identified using the mPD-Q, had significantly worse OKS at baseline and at 2 and 12-months after surgery compared to those with nociceptive pain. The difference in OKS between the neuropathic and nociceptive pain groups in the present study was 7 and 5 points at the 2 and 12-month post-operative time-points, respectively, exceeding the minimal clinically important difference of 5 points on the OKS [6]. Patients with features of neuropathic pain also had a higher risk of moderate to severe long-term pain after arthroplasty, independent of other predictors for poor outcome after surgery.

Similar research includes a study of 17 patients with shoulder impingement who underwent QST and assessment using the PD-Q prior to decompression surgery [21]. Although this study did not show any significant relationship between pre-operative PD-Q score, when used as a continuous measure, and post-operative Oxford Shoulder Score (OSS), the presence of radiating pain or mechanical hyperalgesia preoperatively was significantly associated with a worse OSS post-operatively [21]. A recent study of patients undergoing knee replacement surgery demonstrated that the presence of neuropathic features in the early post-operative period was a predictor of persistent post-operative pain [34]. The results of the present study extend this observation further, and present the opportunity to identify patients with features of neuropathic pain even prior to surgery. This may inform the decision-making process in terms of identifying those with poorer predicted outcomes as well as those who would benefit from adjunctive therapy targeting the features of neuropathic pain pre-, peri-, or post-surgery. Such therapeutic options could include pharmacological (such as duloxetine and amitryptiline) [43] or non-pharmacological approaches (such as education, exercise therapy and cognitive behavioural therapy) [46; 61] [52]. This approach may help to improve outcome following other treatment interventions in the context of osteoarthritis and other musculoskeletal conditions; further data is awaited [27].

The main strengths of this study are the use of prospective, longitudinal data to investigate the relationship between pre-operative neuropathic pain and short and long-term outcome in two independent cohorts, one with a richer description of potential mediators and confounders and a second larger but less well characterized cohort. A variety of clinical, pain and psychological features were measured in the study cohort, enabling the simultaneous assessment of these factors in conjunction with one another. The main limitation is the fact that the primary study cohort did not reach its target recruitment and so was underpowered. The inclusion of data from the larger, validation cohort partly compensates for the relatively small sample size in the study cohort. However, it must be noted that the breadth of data collected in the study cohort is not replicated in the validation cohort.

In summary, this study has shown that the sub-group of patients with knee OA, who have features suggestive of neuropathic pain identified using the m-PDQ, have significantly worse outcome at 2 and 12-months post-operatively compared to those with nociceptive pain. These patients may benefit from increased awareness of their projected outcome to aid informed decision making with respect to surgical intervention. Post-operative outcome may also be improved by the utilisation of targeted therapy in the pre, peri and post-operative periods.

## ACKNOWLEDGEMENTS

The authors would like to thank Alison Turner, Vicky Toghill, Rhea Zambellas, Nicholas Bottomley and William Jackson for their assistance with the EPIONE Study. Anushka Soni was funded by a National Institute for Health Research Fellowship (Doctoral Research Fellowship: DRF-2010-03-131). This paper presents independent research funded by the National Institute for Health Research (NIHR). The views expressed are those of the authors and not necessarily those of the NHS, the NIHR or the Department of Health. Nigel Arden reports personal fees from FLEXION, personal fees from BIOVENTUS, personal fees from MERCK, personal fees from Q-MED, personal fees from ROCHE, personal fees from SMITH & NEPHEW, personal fees from Freshfields, personal fees from BIOIBERICA, outside the submitted work.

Cyrus Cooper reports personal fees from Servier, personal fees from Amgen, personal fees from Eli Lilly, personal fees from Merck, personal fees from Medtronic, personal fees from Novartis, outside the submitted work. Vishvarani Wanigasekera was supported by the NIHR Oxford Biomedical Research Centre. Irene Tracey was supported by the Wellcome Trust. Kirsten Leyland, Amit Kiran, M Kassim Javaid and Andrew Price have nothing to disclose.

